# Spontaneous Pre-encoding Activation of Neural Patterns Predicts Memory

**DOI:** 10.1101/229401

**Authors:** Talya Sadeh, Janice Chen, Yonatan Goshen-Gottstein, Morris Moscovitch

## Abstract

It is well-established that whether information will be remembered or not depends on the extent to which the learning context is reinstated during post-encoding rest and/or at retrieval. It has yet to be determined, however, if the fundamental importance of reinstatement to memory extends to periods of spontaneous neurocognitive activity prior to learning. We thus asked whether memory performance can be predicted by the extent to which spontaneous pre-encoding neural patterns resemble patterns elicited during encoding. Individuals studied and retrieved lists of words while undergoing fMRI-scanning. Multivoxel hippocampal patterns during resting periods prior to encoding resembled hippocampal patterns at encoding most strongly for items that were subsequently remembered. Furthermore, across subjects, the magnitude of similarity correlated with a behavioural measure of episodic recall. The results indicate that the neural scaffold of a memory trace is spontaneously laid even before ever perceiving the to-be-encoded information.

**Significance Statement:** It is well-established that memory performance depends on the degree to which the learning-context is reinstated during post-learning rest or during retrieval. However, does memory also depend on the context *prior* to learning—namely, on processes occurring spontaneously before ever perceiving the to-be-learned information? To answer this question, we scanned participants using fMRI while they were learning and recalling word-lists and, crucially, also during resting periods before each list. Patterns of brain activity in memory-related regions which were elicited spontaneously during these resting periods resembled patterns during learning. Furthermore, the greater this resemblance, the better was memory performance. We demonstrate that memory can be predicted by the degree to which patterns of neural activity prior to learning are reinstated during learning.

## Main Text

Why are certain experiences remembered and others forgotten? The literature has established that memory performance is predominantly dependent on the degree to which the neurocognitive context at encoding is reinstated during post-encoding rest and/or at retrieval (1-5). For instance, using functional magnetic resonance imaging (fMRI), studies in humans have revealed that patterns of brain activity during encoding are similar to patterns elicited during post-encoding rest and during retrieval: the more similar the reinstated pattern to the original one, the better is memory performance (6-15). An open question, however, is whether the fundamental importance of reinstatement to memory is also evident *prior* to encoding. Namely, are spontaneous pre-encoding patterns reestablished during successful encoding and lead to better memory? A positive answer to this question would entail that memory performance can be predicted not only by the degree of post-encoding reinstatement, but also by the degree of pre-encoding *pre-instatement*: that is, the overlap between pre-encoding and encoding patterns. The current study is thus aimed at answering this question, with the hypothesis that the determinants for the mnemonic fate of an experience can be traced back to the time preceding encoding.

Evidence accumulated in recent years has begun to support the viability of this hypothesis. In rodents it has been shown that hippocampal place-cells firing in a particular temporal sequence while an animal is navigating a route also fire spontaneously in the same sequence during a resting period prior to the experience (16-24)^1^. In the absence of this pre-encoding pattern, new patterns must be formed from less activated neurons, thereby reducing their chances of survival in a memory trace (25). Along similar lines, a recent study in humans has found that a “Prototypical Neural Representation” measured several minutes prior to the experimental task predicted understanding of sentences (26). Within the domain of episodic memory, studies investigating processes occurring prior to encoding in humans have found that memory may be enhanced by providing what has been termed an “episodic specificity induction” task, in which participants are briefly trained on recalling details of an event prior to encoding (e.g., 27, 28). Other studies investigated neural activity immediately prior to presentation of each stimulus. These studies found that brain activity (at times, including the hippocampus) predicted memory performance (4, 29-31) and decision-making (32). The findings have typically been interpreted to reflect an anticipatory state, or heightened attention during the pre-stimulus periods. Indeed, oftentimes, pre-stimulus periods were preceded by a cue providing information about the upcoming item.

In contrast to these latter studies, the aim of the current study was not to examine anticipation of an individual experience, or even explicit anticipation at all, but to seek pre-encoding patterns within ongoing spontaneous neural activity that support memory. Based on recent theoretical ideas (24), we hypothesized that pre-existing neural representations provide the framework for successful encoding of new information. Operationally, this entails that, in humans, spontaneous patterns of neural activity during pre-encoding rest are reinstated to support successful encoding. We focused specifically on mnemonic effects in the hippocampus and its related network. We predicted that an item is more likely to be remembered the more similar the neural representation of its memory trace is to a representation spontaneously elicited during pre-encoding rest. Our design thus included 15-20 s resting periods prior to presentation of the study materials (Fig. 1). We used a word-list free recall task, in which participants (n=23) studied and immediately recalled 24 lists of 12 words. During all phases, neural activity was measured using functional magnetic resonance imaging (fMRI). From these data, neural activity patterns for each of the timepoints (TRs) were extracted. This procedure was implemented both for the pre-encoding resting timepoints and for each of the timepoints corresponding to studying a word. A similarity score could then be calculated between each resting timepoint and each study timepoint.

**Fig. 1.**
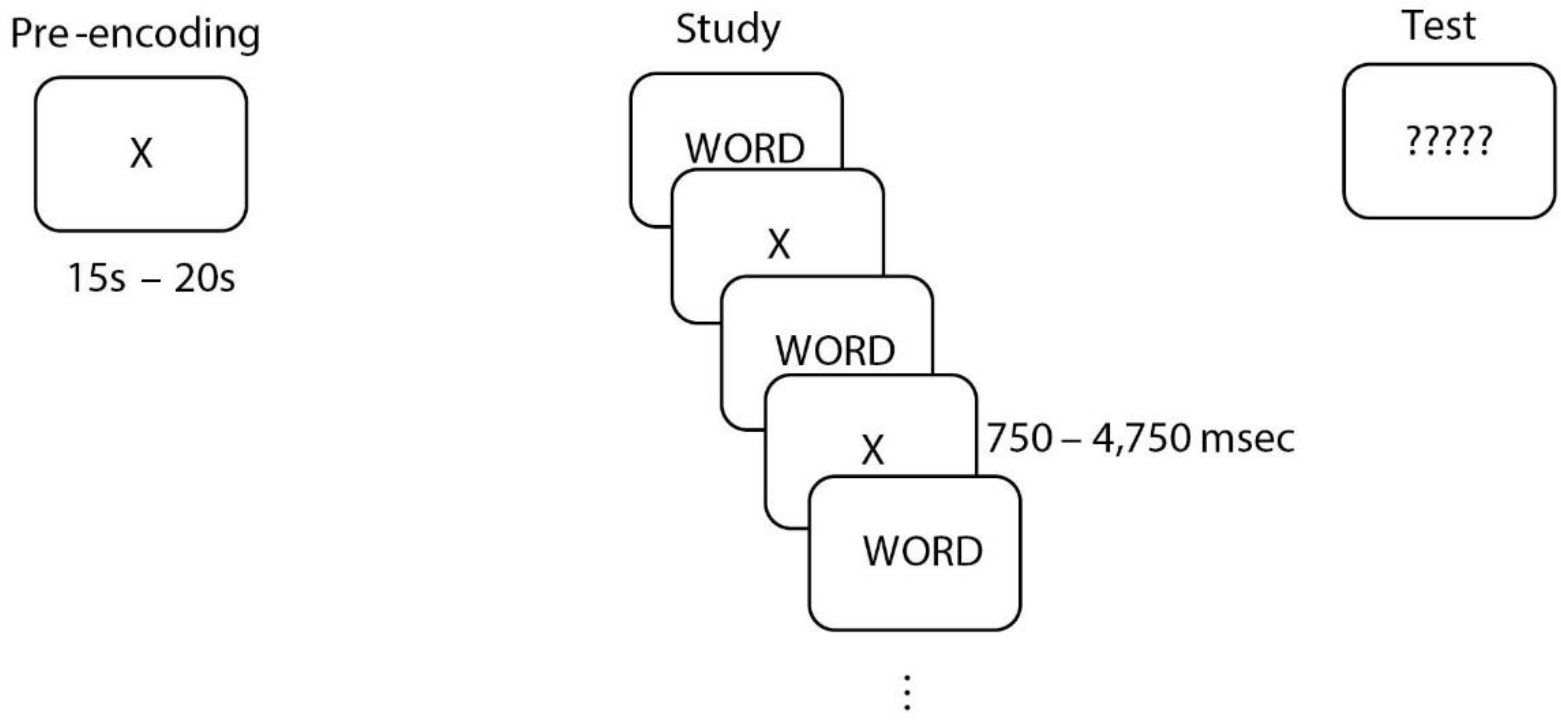
Illustration of the design.

## RESULTS

### Behavioural Results

All reported statistical t-tests are two-tailed. Participants correctly recalled a mean of 6.34 words per list (SEM = 0.36). Of the total number of words recalled 4% were extra-list intrusions and 2.5% were prior-list intrusions, namely words from lists presented previously in the experiment. The prior-list intrusions showed a typical effect of higher probability of intrusions from temporally proximal lists, as compared to more distant lists (e.g., more likely to be a word from the preceding list than from 10 lists prior to the current one; 33). To test the significance of this pattern, we correlated, for each participant, the percentage of intrusions from each of the preceding lists with the distance between the current list and the list from which the intrusion occurred. The average Pearson correlation coefficient was −0.39 and was significantly smaller than zero, as revealed by a one-sample t-test (t(22) = −12.2, p < .001). Recall probability as a function of output position (serial position curve) is presented in Fig. 2. The results show a typical serial position curve with pronounced primacy and recency effects.

**Fig. 2.**
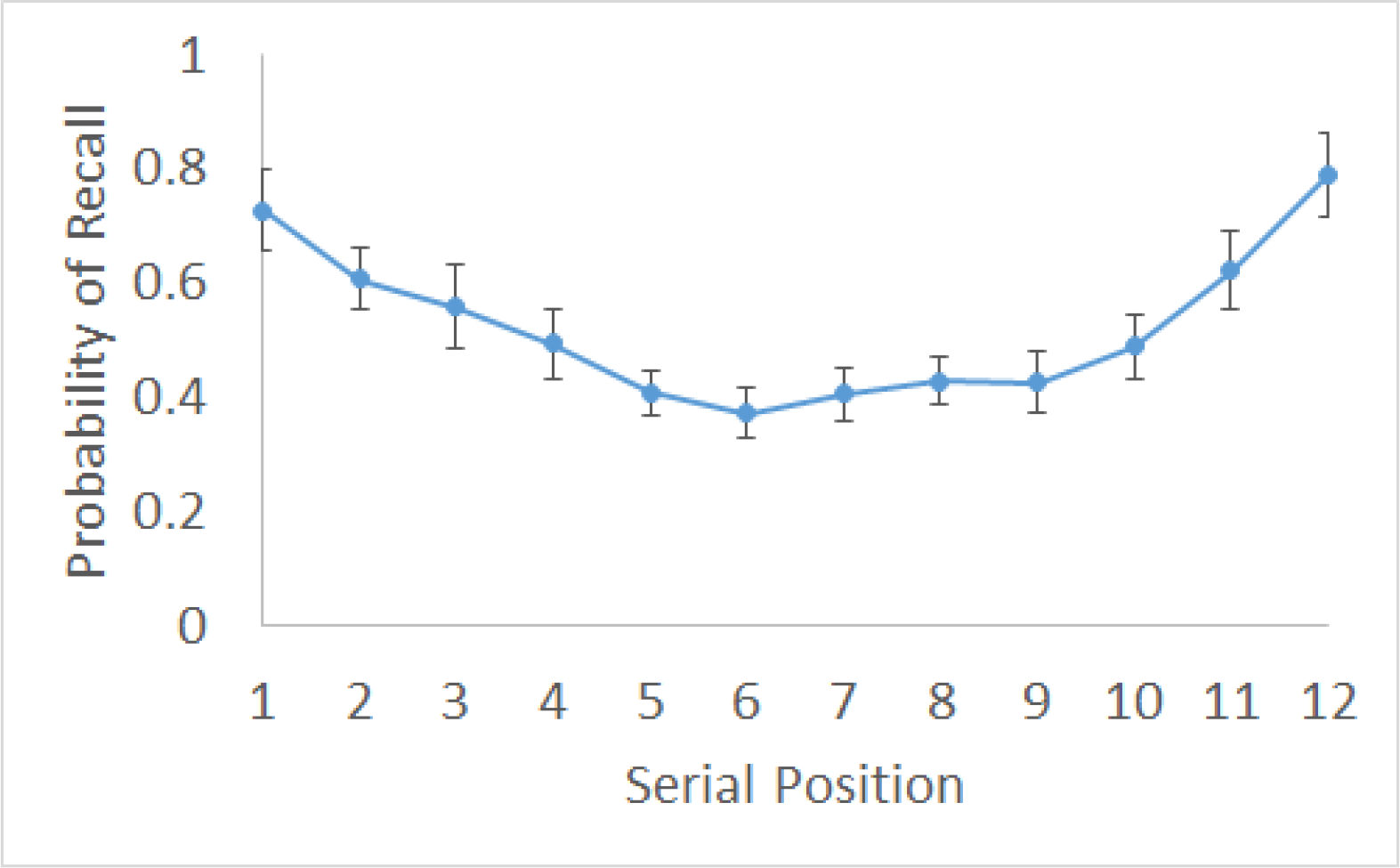
The Serial Position Curve. Probability of recall as a function of study serial position. Error bars reflect 95% confidence intervals for within-subject designs (34).

As illustrated in Figure 3, a typical Temporal Contiguity Effect (35) was observed, demonstrating that the closer two items had been presented at study, the higher their probability of being recalled consecutively. Because we excluded the first three words recalled from each list to eliminate possible effects of working memory, whose span is 3-4 items (36-39), the magnitude of the Temporal Contiguity Effect is comparable to that of delayed free-recall paradigms (35, 40).

**Fig. 3.**
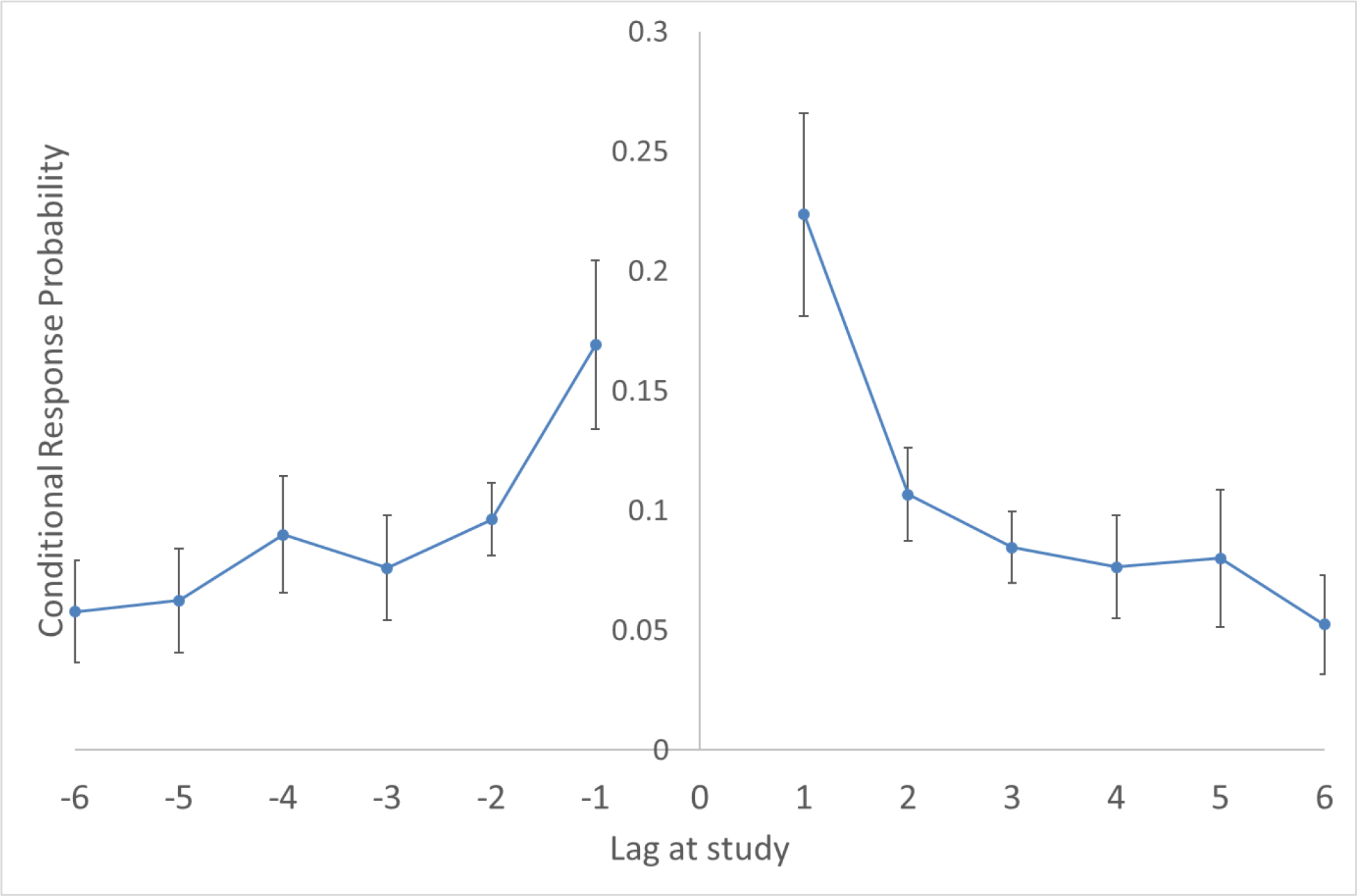
The Temporal Contiguity Effect. The Temporal Contiguity Effect for all but the first three words recalled (which may reflect reliance on working memory rather than episodic memory; see Materials and Methods). Conditional Response Probability (CRP) measures the probability that a transition between two successively-recalled items would be made across a certain lag (35). The probabilities are conditional on the event that a transition of a certain la would yield a studied item that has not already been retrieved. Lag refers to the distance, at study, between the serial-positions of two successively recalled words. Error bars reflect 95% confidence intervals for within-subject designs (34).

### Controlling for distance from pre-encoding between Remembered and Forgotten items

Our analysis of pre-encoding effects was aimed at demonstrating that, as compared to Forgotten items, the neural patterns elicited by Remembered items exhibit enhanced similarity with spontaneous patterns elicited during the pre-encoding phase. It was essential to rule out the possibility that pre-encoding effects are driven by differential distances from pre-encoding between Remembered and Forgotten items: if Remembered items are closer to the pre-encoding phase than Forgotten items, differences between the two conditions could be due to trivial effects of temporal autocorrelation, rather than pre-encoding effects.

For each encoded item, we calculated its distance in TRs from the pre-encoding phase. As might be expected considering the primacy effect, Remembered items were closer to the pre-encoding phase than Forgotten items (Mean Remembered = 7.4 TRs, Mean Forgotten = 8.4 TRs; paired-sample t-test: t(22) = 3.63, p = .001). To control for the effects of distance from encoding, for participants whose Remembered items were, on average, closer to the pre-encoding phase than the Forgotten items (n = 16), we randomly excluded ~25% of the closest Remembered items and ~25% of the farthest Forgotten items. With this exclusion, the distances from pre-encoding were greater (though not significantly; t(22) < 1, p = .9) for Remembered than Forgotten trials (Mean Remembered = 7.64 TRs, Mean Forgotten = 7.62 TRs; for further details see Supplementary Information). All subsequently-reported analyses were conducted on the data controlling for the differences in distances between Remembered and Forgotten items as described above (Mean number of Remembered trials per block = 2.43, S.D. = 0.82; Mean number of Forgotten trials per block = 3.3, S.D. = 1.25). As a further control for distance from pre-encoding, we ran the analysis of pre-encoding effects in the hippocampus excluding the first item in each list.

### Pre-encoding Effects in the Hippocampus

Fisher-transformed Pearson correlation coefficients (41) indexing similarity scores were calculated between the multi-voxel pattern of each pre-encoding TR and of each study trial (see Materials and Methods for further details). For each participant, similarity scores were then averaged across trials for each of the two conditions (Remembered and Forgotten). Figure 4 depicts the results of this analysis for our a-priori region of interest (ROI): the bilateral hippocampi. As predicted, the mean similarity scores in the Remembered condition (Mean Fisher’s Z = 0.42) were greater than in the Forgotten condition (Mean Fisher’s Z = 0.3; paired-sample t-test: t(22) = 2.59, p = .017, Cohen’s d = 0.54). Thus, a significant pre-encoding effect was demonstrated. The additional control analysis excluding the first item from each list (which was the closest to the pre-encoding stage) revealed the same pattern, with greater similarity in the Remembered condition (Mean Fisher’s Z = 0.4) than in the Forgotten condition (Mean Fisher’s Z = 0.26; paired-sample t-test: t(22) = 2.56, p = .018, Cohen’s d = 0.53).

**Fig. 4.**
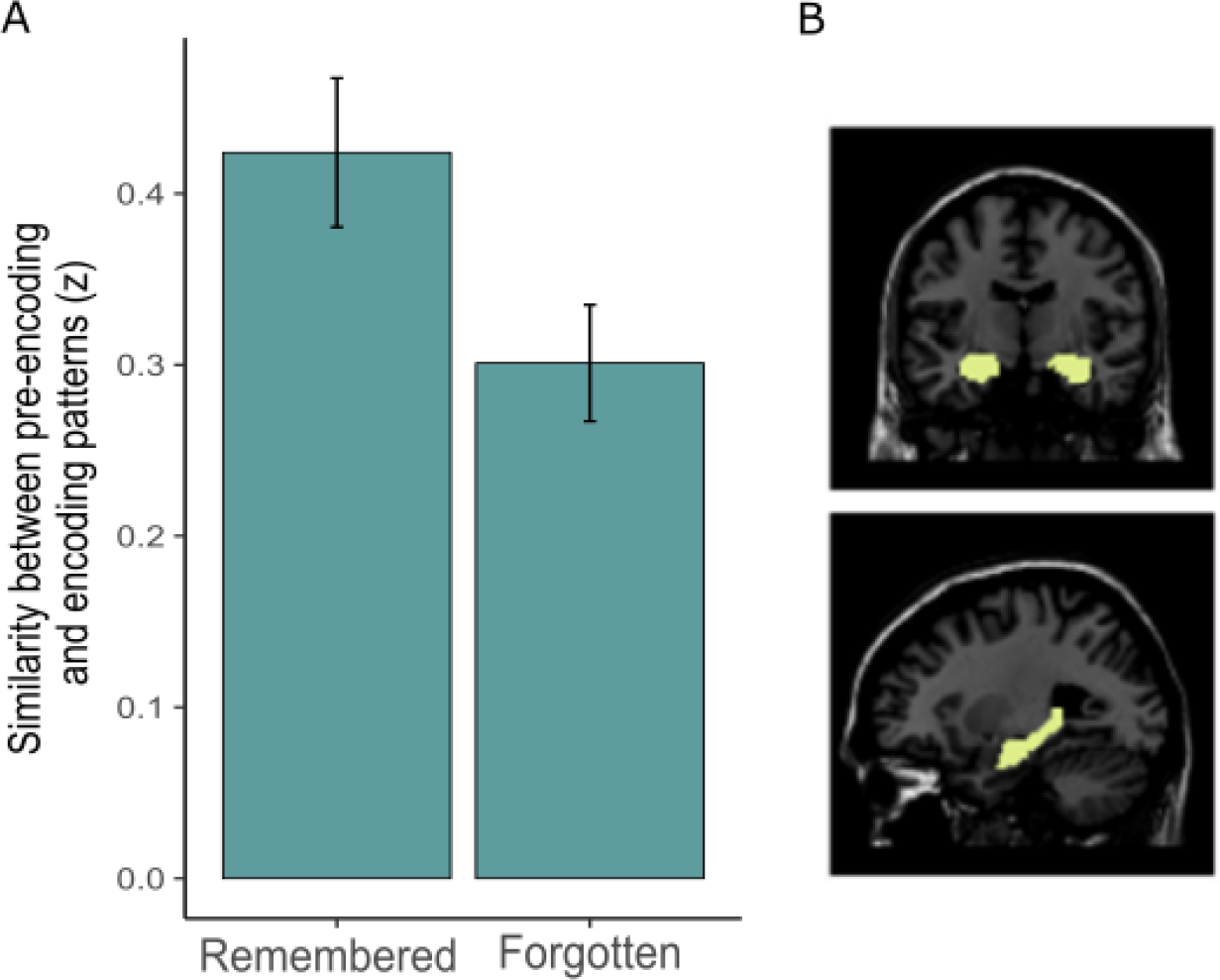
Pre-encoding Effect in the Bilateral Hippocampi. Difference between Remembered and Forgotten words with regard to similarity between pre-encoding and encoding patterns (t(22)= 2.59, p = .017, Cohen’s d = 0.54). Error bars denote standard error of the mean.

### Pre-encoding Effects in the Hippocampal Network

Our a-priori assumption was that pre-encoding reflects the laying down of a scaffold for a memory trace of a future episode. On the neural level, the memory trace consists of an entire hippocampal-neocortical ensemble, in which the hippocampus acts as a pointer to a cortical network representing the perceptual and semantic details of the episode (42, 43). We, therefore, hypothesized that the pre-encoding effect extends to the network of regions which, together with the hippocampus, constitutes the memory trace.

To examine this possibility, we next turned to investigate pre-encoding effects within the network of regions that co-fluctuated with the hippocampus throughout the experiment. To this end, the timecourses for all voxels within the bilateral hippocampi ROI were averaged and used as the seed in a functional connectivity analysis. This analysis aimed to identify voxels whose activity correlates with the hippocampus throughout the experiment. The correlations were calculated separately for each run, then averaged across runs. For each participant, a Hippocampal Network ROI was defined that included all voxels for which the correlation coefficient surpassed a threshold of 0.3^2^. Using the same threshold, a group Hippocampal Network ROI was created by averaging together all of the individual-subjects’ Hippocampal Network ROIs (see Figure 5 for the Group Hippocampal Network ROI). The group-level functional-connectivity analysis revealed a set of regions associated with memory for perceptually-rich experiences (44, 45). These included the parahippocampal gyrus, occipital-temporal regions, the posterior cingulate cortex, lingual gyrus and precuneus. In addition, large clusters were detected in the thalamus, the striatum and regions in the cerebellum.

**Fig. 5.**
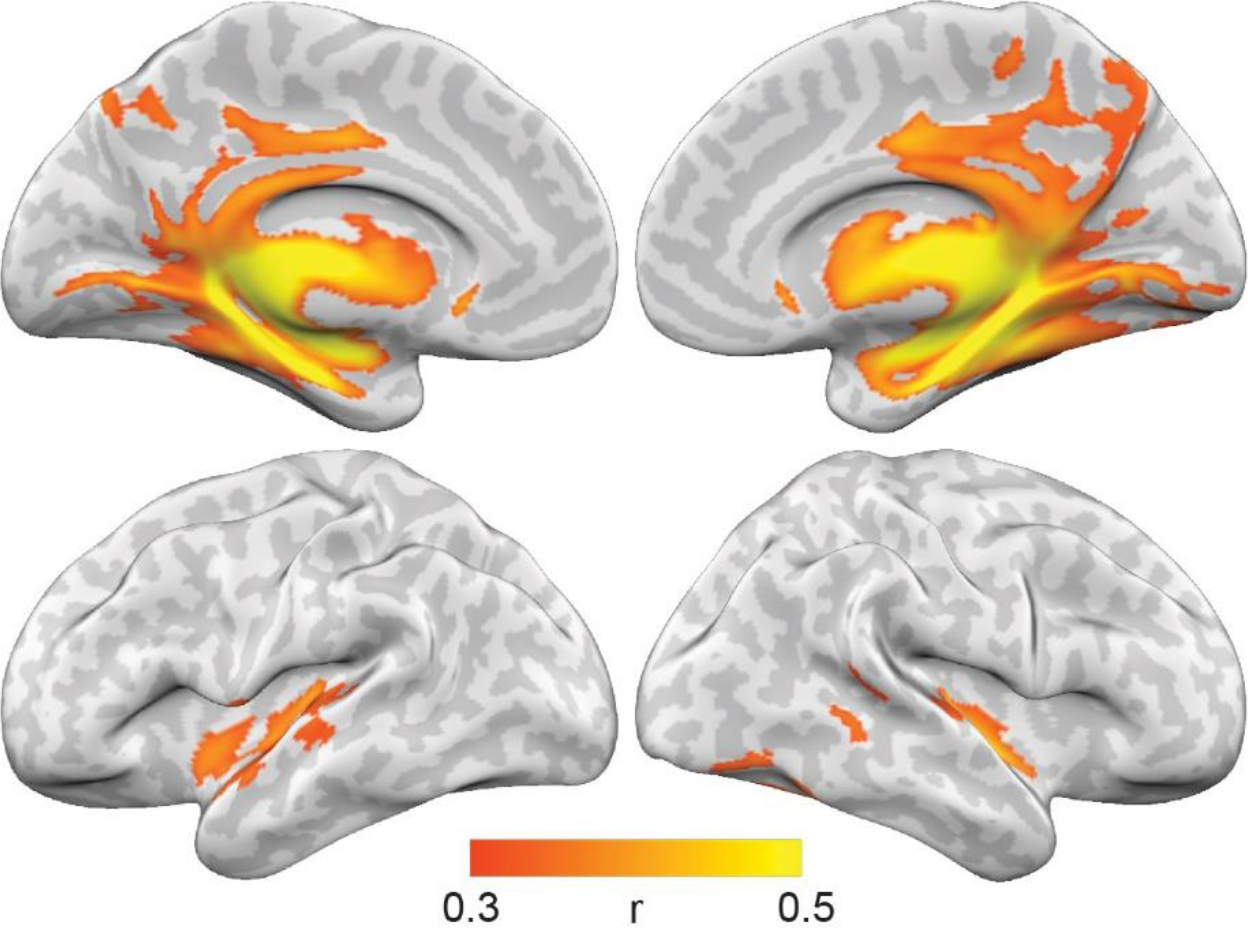
Group Hippocampal Network ROI. Regions co-fluctuating (functionally connected) with the hippocampus during the experiment (voxels for which the mean correlation coefficient surpassed a threshold of 0.3, averaged across participants).

The analysis comparing pre-encoding effects between the Remembered and Forgotten conditions was repeated for these ROIs. As in the analysis of the hippocampus, here, too, the mean similarity score in the Remembered condition (Mean Fisher’s Z = 0.19) was significantly greater than in the Forgotten condition (Mean Fisher’s Z = 0.096), as revealed in a paired-sample t-test (t(22) = 3.32, p = .003, Cohen’s d = 0.69). In the Group Hippocampal Network ROI, the effect was also significant (Mean Fisher’s Z for Remembered = 0.14; Mean Fisher’s Z for Forgotten = 0.038; t(22) = 2.97, p = .007, Cohen’s d = 0.62).

Importantly, though these analyses reveal that the pre-encoding effect extends to brain regions which are functionally-coupled with the hippocampus, it is unlikely to be a whole-brain effect, as revealed by a follow-up analysis. This analysis explored 90 individual AAL ROIs spanning the entire brain and found that only a subset of 26 regions—20 of which overlapped with the Hippocampal Network ROI—showed a significant effect at p < .05 (for the full list see Table S1). Furthermore, none of these regions passed correction for multiple comparisons (q(FDR) < .05 (46)).

### Across-subject Correlation between Pre-encoding/Encoding Overlap and the Context Reinstatement

Our a-priori hypothesis was that memory depends on the extent to which contextual associations are *pre-instated* prior to encoding. That is, during encoding, the contextual associations evoked by a studied item reinstate thoughts or associations evoked spontaneously during pre-encoding rest. A natural prediction that follows this hypothesis is that the magnitude of pre-encoding/encoding overlap would be associated with reliance on contextual processing. To test this prediction, we capitalized on the Temporal Contiguity Effect (35, 47)—one of the most robust behavioural findings establishing the role of context in driving recall (see Fig. 3). The TCE refers to the increased probability of sequentially recalling two items that were studied in close temporal contiguity. The effect is thought to arise from the largely overlapping temporal (and neural) contexts shared by neighbouring items, and is a hallmark of episodic recall (48, 49).

We thus examined the across-subject correlation between (1) the magnitude of overlap between pre-encoding and encoding representations and (2) the TCE per participant. The magnitude of pre-encoding/encoding overlap per participant was indexed by the mean correlation coefficients between pre-encoding trials and subsequent encoding trials. With regard to the behavioural measure, for each participant, a temporal factor score was calculated. This score represents the tendency of a participant to successively retrieve items in short temporal lags, or to rely on temporal context at recall (see “Behavioural Analysis”). A significant positive correlation was found both for the bilateral hippocampus and for the Hippocampal Network ROI: the greater the pre-encoding/encoding overlap, the stronger the reliance on temporal context (for bilateral hippocampus: r = .56, p = .006; for Hippocampal Network ROI: r = .56, p = .006). Results for the bilateral hippocampus and the Hippocampal Network are illustrated in Figure 6.

**Fig. 6.**
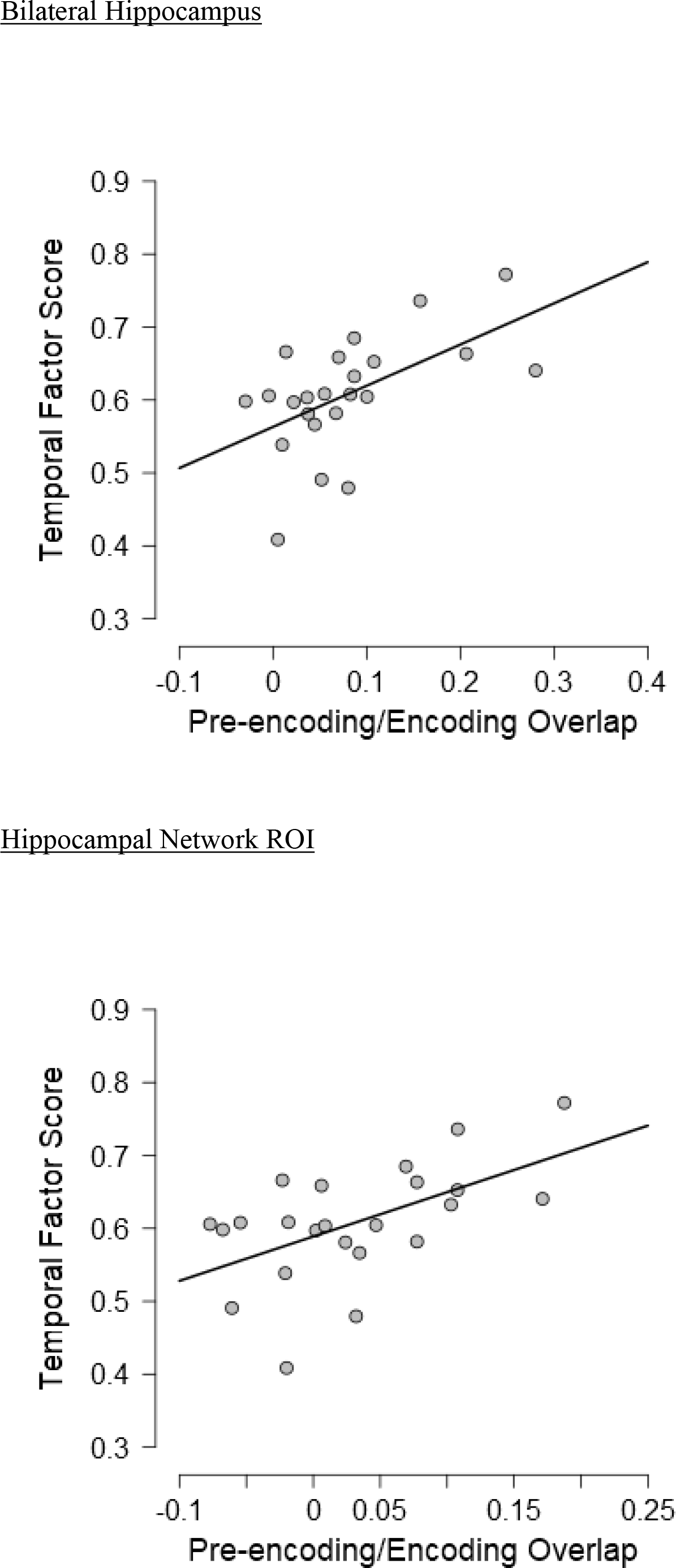
Correlation between pre-encoding/encoding overlap and the Temporal Contiguity Effect.

### Waxing and waning of a general encoding state?

We maintain that the pre-encoding effects demonstrated here reflect correspondence between specific pre-encoding neural patterns and neural patterns of subsequently-experienced events which form the memory trace. However, an alternative interpretation of our findings is that the pre-encoding effects are a result of a general encoding state (or even an attentional state; see Discussion) that waxes and wanes over time. According to this interpretation, the pre-encoding effects reflect a general similarity between pre-encoding and encoding neural states that is not item-specific.

To examine this possibility, we ran a sham analysis in which we aggregated across all lists to create two correlation matrices per subject: one including correlation coefficients between all pre-encoding trials (across all lists) and all subsequently-remembered encoding trials, and one between all pre-encoding trials and all subsequently-forgotten trials. An average correlation coefficient was then calculated (and Fisher-transformed) for each of these two matrices. A sham pre-encoding effect was defined as the difference between these two averages—namely, between the subsequently-remembered average Fisher-transformed correlation coefficient and the subsequently-forgotten average Fisher-transformed correlation coefficient. If the alternative, “encoding state”, interpretation is true, it is expected that the sham pre-encoding effects would be of a similar magnitude as the original pre-encoding effects. Results of the sham analysis argue against this interpretation. For both the hippocampus and the Hippocampal Network ROI, the original pre-encoding effects were significantly greater than the sham pre-encoding effects (for the hippocampus: t(22) = 2.33, p = .029, Cohen’s d = 0.49; for the Hippocampal Network ROI: t(22) = 2.72, p = .012, Cohen’s d = 0.57; Figure 7). This indicates that the patterns of remembered items are more similar to the pre-encoding phase which immediately preceded them as compared to the pre-encoding phases of the other lists.

**Fig. 7.**
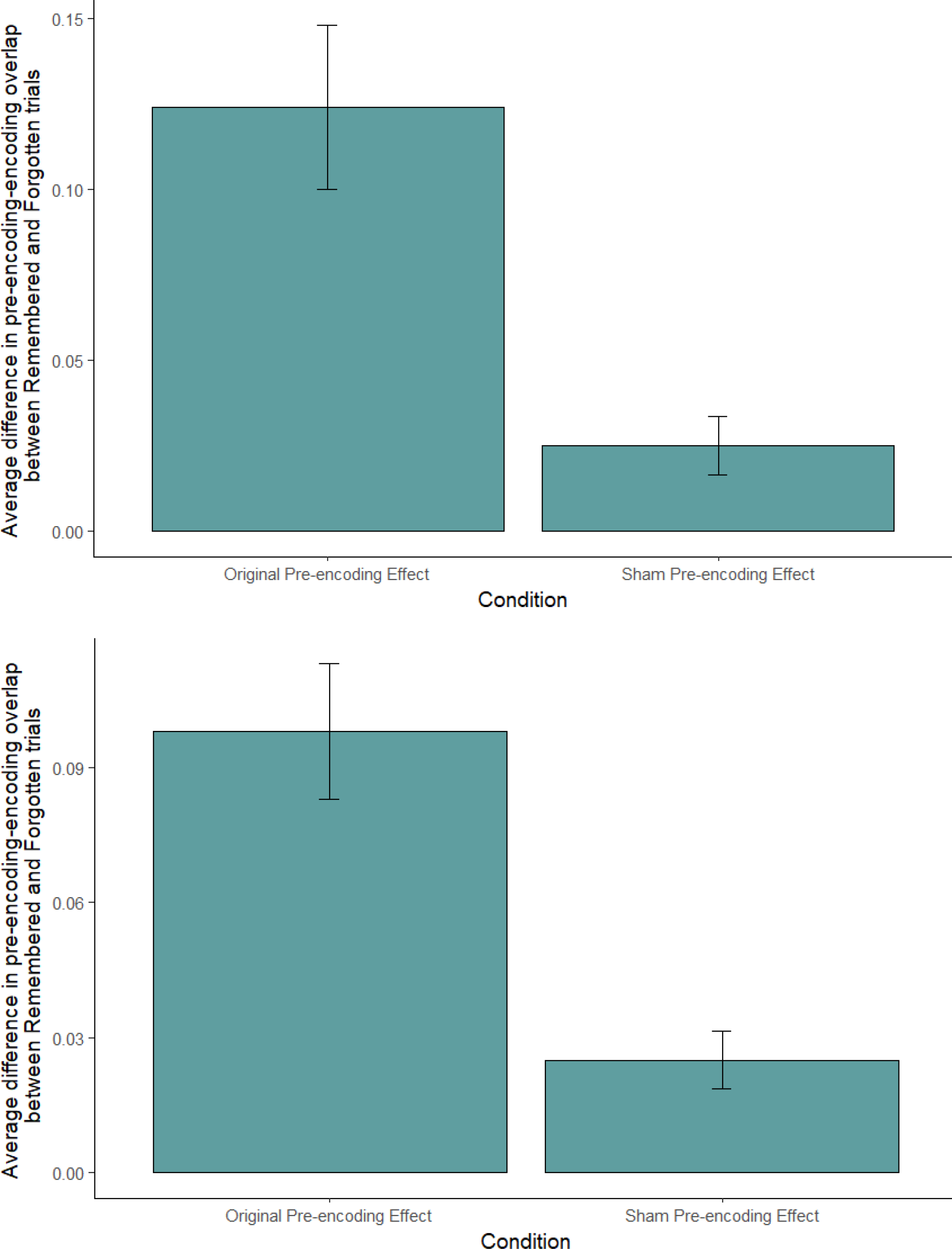
Comparison of Original and Sham Pre-encoding Effects. Pre-encoding effects are the difference between Remembered and Forgotten trials with regard to similarity between pre-encoding and encoding patterns. The original pre-encoding effect refers to similarity between the pre-encoding trials preceding a certain list and the encoding trials of that specific list. The sham pre-encoding effect refers to similarity between all pre-encoding trials (across all lists) and all encoding trials. The top panel depicts pre-encoding effects in the bilateral hippocampus and the bottom panel depicts pre-encoding effects in the Hippocampal Network ROI. Error bars denote standard error of the mean.

## Discussion

Our results demonstrate that spontaneous neural patterns elicited during an ongoing pre-encoding resting period are reinstated during successful encoding of individual items. The pre-encoding effects we found are not merely a result of temporal autocorrelation, as revealed by our controls for temporal distance. As predicted, these effects were found in item-specific spatial activity patterns of the hippocampus, as well as in its functionally-coupled network. Furthermore, we found that individual differences in the magnitude of pre-encoding/encoding overlap correlated with reinstatement of context at retrieval, as indexed by the temporal contiguity effect.

The Temporal Contiguity Effect refers to the increased probability of sequentially recalling two items that were studied in close temporal contiguity. The effect is thought to arise from the largely overlapping temporal (and neural) contexts shared by neighbouring items and its reinstatement during recall. What can be conceptualized as a neural manifestation of this effect has been proposed in the rodent literature in the theoretical framework of Memory Allocation (25). According to the notion of Memory Allocation, neurons with intrinsically high excitability are more likely to be allocated to a memory trace of an item. These neurons are also more likely to be included in the memory traces of items studied adjacently to the given item. Consequently, neighbouring items at study are also more likely to cue each other during test and thus be clustered together at retrieval. This clustering phenomenon is exactly that which is documented in the context of the Temporal Contiguity Effect.

Are the pre-encoding patterns merely random variations in neural firing that are fortuitously co-opted by neural events at encoding? If so, does the pre-encoding effect arise from a general encoding or attentional state (50) that waxes and wanes during pre-encoding and encoding? If this interpretation is true, the overlap between pre-encoding and encoding patterns reflects points in time in which the mnemonic system is in a preferred, or good, encoding and/or attentional state. If so, we would expect encoding patterns to overlap not only with patterns elicited during the preceding pre-encoding block, but with patterns elicited during all resting blocks in the experiment. Our sham analysis reveals that this is not the case—namely, that our results cannot be fully accounted for by the notion of a general encoding or attentional state. Rather, our results are at least partially (if not fully) driven by spontaneous pre-instatement of item-specific neural patterns.

While the particular states or thoughts represented by the pre-encoding neural patterns are not amenable to direct investigation, it is possible that they reflect, at least in part, episodic thoughts which are idiosyncratically associated with the to-be-studied items. This idea does not, of-course, entail any form of pre-cognition (51). Namely, we do not claim that during pre-encoding rest participants spontaneously thought of the exact same words that subsequently appeared during the study phase. Rather, it is possible that spontaneous idiosyncratic thoughts during pre-encoding became associated with (at least some of) the study words and constituted part of their memory traces, perhaps due to shared contextual features between the spontaneous thoughts and the memory traces of the study words. Such spontaneous thoughts and associations are constantly evoked during rest—a finding well-established in the literature regarding the brain’s default mode at rest (52-54). These thoughts and associations are the cognitive correlate of the presumed pre-existing neural representations which support encoding of new information and give rise to the pre-encoding effects. Although it would be interesting, and informative, if there were such cognitive correlates, their presence is not crucial for our results to be valid.

In our paradigm, pre-encoding resting periods most frequently followed periods of recall of words from the previous lists. This raises the possibility that the pre-encoding patterns were influenced by words from the previous lists. Thus, the pre-encoding patterns may have reflected reinstatement of the contextual associations evoked by items from previous lists. Consistent with the current findings within the hippocampus, effects of replay of individual items within the medial temporal lobe have been recently reported (6). The “bleeding” of contextual representations from previous lists to encoding of a given list has also been previously shown on the behavioural level (e.g., 33, 55-57), and was evident in the current study, where prior-list intrusions were more likely to be from temporally proximal lists, as compared to more distant lists.

All pre-encoding effects reported so far were found both for the a-priori bilateral hippocampal ROI and for the network of regions functionally coupled with the hippocampus, (referred to in the Results sections as “Hippocampal Network ROIs”). This latter result held both when the functional network was defined per subject and when defined at the group level. An additional analysis (see Supplementary Information) sought to identify whether the network results reflect only patterns within individual ROIs (or subsets of the network) or also a global pattern of amplitudes across the different ROIs in the network—namely, a network effect. The analysis, which we term the “Global-pattern ROI”, also revealed a significant pre-encoding effect. This finding suggests that the current pre-encoding effects may be driven in part by a global pattern of amplitudes (namely, by network-wide effects).

Demonstrating the existence of a mechanism of pre-encoding provides a crucial contribution to our understanding of why certain experiences are remembered and others are forgotten. The mnemonic fate of studied-items greatly depends on the extent to which spontaneous neural representations, within an ongoing pre-encoding period, are reinstated during encoding. Because the individual’s neural state at the resting time preceding encoding likely differs from one occasion to the next—and correspondingly so do the idiosyncratic thoughts/autobiographical associations which are prominent during rest (52, 53)—so will the mnemonic outcome of studied items be different, even if all external stimuli and conditions are kept the same in all occasions.

Whatever the specific nature of pre-encoding patterns might be, our findings illustrate that they play a crucial role in memory for subsequently-presented information. With that we provide a novel demonstration in humans that the determinants for the mnemonic fate of an experience can be traced back to spontaneously-elicited neural patterns prior to the experience. Thereby, we establish that pre-encoding constitutes a fundamental aspect of the neurocognitive basis of human memory. These findings have important implications for memory-interventions in healthy and in memory-impaired people from early development to aging, by extending the focus of such interventions to processes occurring prior to presentation of memoranda.

## Methods

### Participants

Participants were 28 neurologically-intact native Hebrew speakers (17 women), right-handed and with normal or corrected-to-normal vision. Data from four participants were excluded due to excessive motion inside the scanner (over 4 mm). Data from an additional participant were excluded due to low performance on the behavioural memory task (mean number of recalled items was over two standard deviations below the group average). All reported analyses thus include data of 23 participants (15 women; ages 21-32 years, mean = 24.5). Participants gave their informed consent prior to the experiment and were compensated for their time monetarily or with course credit. All experimental procedures were approved by the Tel-Aviv Medical Center’s Clinical Investigation committee.

### Materials

The stimuli consisted of 336 Hebrew nouns. All nouns were 5-6 letters long (mean = 5.46). The nouns were divided into 28 lists of 12 words; 24 lists were used for the experiment and 4 lists for the practice phases (see below).

### Behavioral Procedure

Prior to entering the scanner, participants were given detailed instructions and two practice lists of the free-recall task, which included overt pronunciation of the encoded words. A primary goal of this practice session was to ensure that participants verbalized the words clearly yet softly enough to avoid head motion^3^. Two additional practice lists were presented within the scanner.

The experiment consisted of four identical free-recall runs, each lasting 7:25 minutes. The set of words was randomly divided into four, with a quarter of the words (i.e., 72 words, divided into six lists of 12 words each) assigned to each run. The order of the runs was counterbalanced across participants. In addition, three runs of a semantic fluency task were interleaved in between the free-recall runs. The purpose of the semantic fluency task was extraneous to the current endeavor, and therefore we do not discuss it further.

In each of the four runs, six lists of 12 words each were presented for study followed by a free-recall test. The order in which the words were presented was random. Presentation of three of the six lists (randomly selected) was preceded by a fixation block, which we term the “Pre-encoding Phase”. During this phase, participants were instructed to rest while fixating on a cross in the middle of the screen. The durations of the pre-encoding phases were 15, 17.5 and 20 s and their order was counterbalanced across runs. Only lists which included a pre-encoding phase were included in the current analyses.

Presentation of each of the study lists was preceded by a 2.5 s display in which the word PREPARE appeared at the center of the screen, signaling participants to prepare for the upcoming list. The study phase then began, with each of the 12 words in the list presented sequentially for 1750 msec in the center of the screen followed by a 750-4,750 msec fixation cross. For four of the six lists (whose order was randomly assigned and counterbalanced across sessions), a fixation trial of 0.5 to 8.25 s was presented at the offset of the study phase. The aim of these trials was to jitter the beginning of the recall phase with regard to the TR. The recall phase then began, with presentation of five question marks at the center of the screen signaling participants to start recalling. Recall was executed by overtly pronouncing as many words as possible from the last list presented, in any order, until the cue preparing them for the next list appeared on the screen. The recall phase lasted 22.5 s. Verbal responses were digitally recorded using Audacity software (http://audacity.sourceforge.net). As an incentive to enhance performance, participants were told that they would be awarded monetary prizes (comparable to $200) if they reached the highest scores in the experiment. Figure 1 illustrates the experimental design for a single list which is preceded by a pre-encoding phase.

### Imaging Procedure

Participants were scanned on a GE 3T Signa Horizon LX 9.1 echo speed scanner (Milwaukee, WI). During each of the runs, whole-brain T2*-weighted EPI functional images were acquired (TR=2500 msec, 20 cm FOV, 64x64 matrix, Flip Angle=85, TE=35, 44 coronal slices perpendicular to the hippocampal axis, 3 mm thickness with 0.7 gap, sequential acquisition). In each run, 178 volumes were acquired. Four additional volumes were acquired at the beginning of each run to allow for T1 equilibration (and were excluded from the analysis).

### Data Analysis

#### Behavioural Analysis

The behavioural recall data were transcribed manually by research assistants. Based on the transcription, the study items were classified into the following categories: (1) Words subsequently recalled at the first three output positions. These presumably reflect retrieval from working memory, whose span is 3-4 items (37, 39), which was not the focus of the current study. Crucially, these items were not included in any of the analyses, except the one examining the serial position curve. (2) Words subsequently recalled correctly, from all output positions but the first three (“Remembered”); (3) Words which were not subsequently-recalled (“Forgotten”).

The Temporal Contiguity Effect was examined using scripts from http://memory.psych.upenn.edu/Software. To reiterate, the Temporal Contiguity Effect refers to the phenomenon whereby the smaller the absolute temporal lag between two items at study, the higher the probability that these two items will be recalled consecutively. We examined conditional-response probabilities (CRPs): the probability of making transitions at a certain lag conditional on this lag being available (35). For the analysis examining across-subject correlation with pre-encoding magnitude, temporal-factor scores were calculated for each participant (58). The temporal factor score is a measure of the tendency of a participant to successively retrieve items with short temporal lags (namely, which appeared close to each other at encoding), with a score of 0.5 indicating no effect of temporal contiguity.

#### Pre-processing

Imaging data were preprocessed and analyzed using SPM8 (Wellcome Department of Cognitive Neurology, London). A slice-timing correction to the first slice was performed followed by realignment of the images to correct for subject movement. Next, data were spatially normalized to an EPI template based upon the MNI305 stereotactic space (59). The images were resampled into 2 mm cubic voxels and spatially smoothed with an 8 mm FWHM isotropic Gaussian kernel. Finally, images were resliced to be aligned to the anatomical ROIs.

#### Pattern Similarity Analysis

Effects of pre-encoding were investigated using a method introduced by Staresina et al. (6) to study reactivation of individual items. The pre-encoding phases were broken down to pre-encoding trials, each trial corresponding to single timepoint, or TR. Likewise, the study phase was broken down by TRs. Since study trials were not temporally aligned with the TRs, the onset of each study trial was defined as that of the closest TR. TRs corresponding to trials whose onsets overlapped those of another trial were excluded from the analyses. As suggested in a recent methodological analysis (60), all trials were shifted forward in time by two TRs (5 seconds), since the peak BOLD response is ~5 seconds after the onset of each trial. For each trial, a multi-voxel pattern of activity within regions of interest was extracted from the BOLD data. Data were detrended to remove linear drifts. A similarity score, indexed by Pearson correlation, was then calculated between the pattern of each pre-encoding TR and of each study trial. A Fisher transformation was applied to the correlation coefficients, aiming to make their sampling distribution approach that of the normal distribution, and the results were divided by the coefficientsʼ standard deviations (1/√n − 3) (41).

#### Regions of Interest Definition

Masks of 90 cortical and subcortical regions spanning the brain (excluding the cerebellum) were obtained using the AAL (Automated Anatomical Labeling) atlas (61).

**REFERENCES**

## Acknowledgments

T.S. is grateful to the Azrieli Foundation for the award of an Azrieli Fellowship. We thank Richard Henson and Bradley Buchsbaum for their thoughtful and insightful comments on this manuscript.

## Footnotes

1 It has recently been suggested that this finding may only be obtained if the animal is already familiar with the spatial environment or the goal, but the path to the goal is novel.

2 The connectivity analysis is not redundant with the one finding a Pre-encoding effect in the hippocampus. While the hippocampal Pre-encoding effect concerns spatial correlation and is calculated on an item-by-item basis, the connectivity analysis concerns temporal correlation and is calculated across the whole session.

3 We were able to record overt responses during the fMRI scanning sessions by using adaptive noise cancelling microphone and headphones (FOMRI-III; Optoacoustics, Israel; see also Sadeh et al., 2011).

